# What is unique about the human eye? Comparative image analysis on the external eye morphology of human and nonhuman great apes

**DOI:** 10.1101/2021.09.21.461196

**Authors:** Fumihiro Kano, Takeshi Furuichi, Chie Hashimoto, Christopher Krupenye, Jesse G. Leinwand, Lydia M. Hopper, Christopher F. Martin, Ryoma Otsuka, Tomoyuki Tajima

**Affiliations:** Center for the Advanced Study of Collective Behavior (CASCB), University of Konstanz, Konstanz, 78464, Germany; Max-Planck Institute of Animal Behavior, Radolfzell am Bodensee, 78315, Germany; Kumamoto Sanctuary, Wildlife Research Center, Kyoto University, Uki, Kumamoto, 8693201, Japan; Primate Research Institute, Kyoto University, Inuyama, Aichi, 4848506, Japan; Department of Psychological & Brain Sciences, Johns Hopkins University, Baltimore, MD, 21218, USA; Department of Psychology, Durham University, Durham, DH1 3LE, UK; Lester E. Fisher Center for the Study and Conservation of Great Apes, Lincoln Park Zoo, Chicago, IL 60614, USA; Department of Molecular and Comparative Pathobiology, Johns Hopkins University School of Medicine, Baltimore, MD, 21205, USA; Indianapolis Zoo, Indianapolis, IN 46222, USA; Graduate School of Asian and African Area Studies (ASAFAS), Kyoto University, Kyoto, 6068304, Japan; Wildlife Research Center, Kyoto University, Kyoto, 6068203, Japan; Unit of Synergetic Studies for Space, Kyoto University, Kyoto, 6068306, Japan

**Keywords:** eye color, communication, comparative analysis, human evolution, great ape, sclera, gaze detection

## Abstract

The gaze-signaling hypothesis and the related cooperative-eye hypothesis posit that humans have evolved special external eye morphology, including exposed white sclera (the white of the eye), to enhance the visibility of eye-gaze direction and thereby facilitate conspecific communication through joint-attentional interaction and ostensive communication. However, recent quantitative studies questioned these hypotheses based on new findings that humans are not necessarily unique in certain eye features compared to other great ape species. Therefore, there is currently a heated debate on whether external eye features of humans are distinguished from those of other apes and how such distinguished features contribute to the visibility of eye-gaze direction. This study leveraged updated image analysis techniques to test the uniqueness of human eye features in facial images of great apes. Although many eye features were similar between humans and other species, a key difference was that humans have uniformly white sclera which creates clear visibility of both eye outline and iris –the two essential features contributing to the visibility of eye-gaze direction. We then tested the robustness of the visibility of these features against visual noises such as darkening and distancing and found that both eye features remain detectable in the human eye, while eye outline becomes barely detectable in other species under these visually challenging conditions. Overall, we identified that humans have distinguished external eye morphology among other great apes, which ensures robustness of eye-gaze signal against various visual conditions. Our results support and also critically update the central premises of the gaze-signaling hypothesis.

## 1. Introduction

The gaze-signaling hypothesis proposed by Kobayashi and Kohshima (1997, 2001) posits that the unpigmented exposed sclera of human eyes (i.e., the white of the eye) enhances the visibility of eye-gaze orientation, thereby enabling the eye-gaze signal to function as a powerful communicative device in humans. Their complementary gaze-camouflaging hypothesis exploited the opposite logic for nonhuman primates, proposing that the pigmented exposed sclera of nonhuman primates conceals the visibility of their eye-gaze orientation, particularly the direct gaze, to predators or dominant conspecifics. These hypotheses derive from Kobayashi and Kohshima’s findings that, 1) among 88 systematically studied primate species, exposed white sclera was a unique feature of the human eye; 2) in humans, the sclera was more widely exposed than in other species; and 3) human eyes were horizontally more elongated than those of other primates. The related cooperative-eye hypothesis (Tomasello, Hare, Lehmann, & Call, 2007) extended the gaze-signaling hypothesis to propose that humans have evolved such unique eye morphology to enhance joint attentional and communicative interactions among conspecifics, critical ingredients for human cooperation. Other theorists also noted that the readability of human eye-gaze orientation is critical in other hallmark human group activities, such as cultural transmission and language learning (Csibra, 2010; Csibra & Gergely, 2009).

Despite the widespread popularity of the gaze-signaling hypotheses in the literature, this hypothesis has been severely challenged by recent quantitative morphological studies based on new findings that the human eye is not necessarily unique compared to the eye of other great ape species in terms of shape and color (Caspar, Biggemann, Geissmann, & Begall, 2021; Mayhew & Gómez, 2015; Perea-García, Kret, Monteiro, & Hobaiter, 2019). As one of the first follow-up studies, Mayhew and Gómez (2015) collected a larger sample of images of gorillas and humans than Kobayashi and Kohshima (2001) and found that, although human eyes are indeed more elongated than gorilla eyes, the sclera is exposed to a similar degree in both species, especially in averted eyes. Caspar et al. (2021) recently replicated this same result, further questioning the proposed uniqueness of human eye morphology. However, these previous studies only measured sclera exposedness in the horizontal dimension (called the Sclera Size Index in Kobayashi and Kohshima, 2001). Kaplan and Rogers (2002) pointed out that the degree of sclera exposedness should be examined two-dimensionally (i.e., the area of sclera that is exposed), because primates move their eyes both horizontally and vertically. In fact, one of Kobayashi and Kohshimas’ (2001) findings showed that primates with more arboreal living styles move their eyes more vertically than those species with more terrestrial living styles (e.g., nonhuman apes as compared to humans). Thus, it remains unclear whether the sclera is more widely exposed (area-wise) in the human eye compared to the eye of other ape species.

These follow-up studies also questioned the uniqueness of human eye colors. Mayhew and Gómez (2015) examined both individual and species differences in sclera pigmentation using a large sample of gorillas and humans and found that while all of the examined human individuals had all-white sclera (depigmented all the way from the iris edge to the eye corners), there were substantial individual differences in the extent to which the exposed sclera was depigmented particularly among western lowland gorillas. More recently, Perea-García et al. (2019) measured the contrast between the highest and lowest lightness values within an eye image using a large sample of bonobos, chimpanzees, and humans and Casper (2021) extended this effort to include all great and lesser ape species. This work measured the contrast between iris and sclera: high lightness contrast indicates either high lightness in sclera and low lightness in the iris or the opposite. Among great apes, the former pattern of eye color was observed in humans, lowland gorillas, and Sumatran orangutans, whereas the opposite pattern was observed in chimpanzees and mountain gorillas (Figure 1). Overall, lightness contrast did not generally differ between humans and nonhuman apes, irrespective of the pattern of iris-sclera color. The authors thus suggested that eye-gaze is as conspicuous in nonhuman apes as in humans.

**Figure 1.**
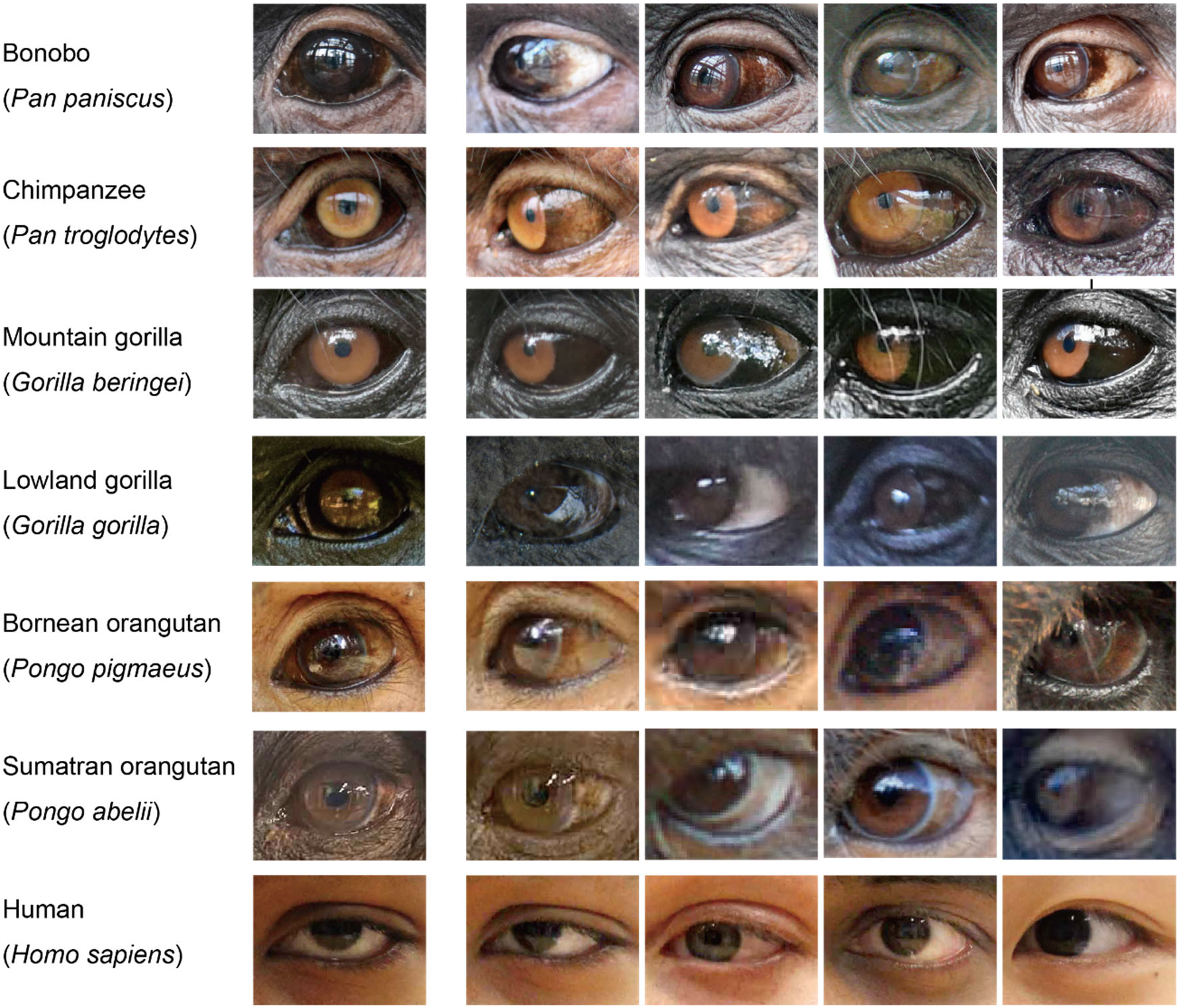
Examples of eye images from great ape species. In the first and second columns, direct and averted eyes of the same individuals are presented. Permission was obtained for the publication of human images from the authors of the public database, The Columbia Gaze Data Set (Smith et al., 2013).

This recent work on iris and sclera color and contrast cast doubt on the gaze-camouflaging hypothesis (concealment of direct gaze). However, it remains unclear how these findings relate to the gaze-signaling hypothesis (advertisement of gaze directions) because, as Kobayashi and Kohshima (2001) pointed out, the visibility of eye-gaze orientation critically depends on the visibility of iris as well as eye outline. Namely, while a strong iris-sclera color contrast ensures the visibility of eye *per se* in the face, clear visibility of both eye outline and iris is essential to ensure the visibility of eye-gaze direction (see Figure 2). It thus remains unclear how humans and nonhuman apes compare in their visibility of these two eye features. Moreover, the eye-color coding method employed by Perea-García et al. (2019) has been critisized by more recent studies (Caspar et al., 2021; Mearing & Koops, 2021). We also note that their method (measuring a ratio between the highest and lowest lightness values in the eye) may be too simplistic to capture the complex color patterns of great ape eyes; specifically, graded and patchy pattens of sclera and the colorfulness of the iris (see Figure 1).

**Figure 2.**
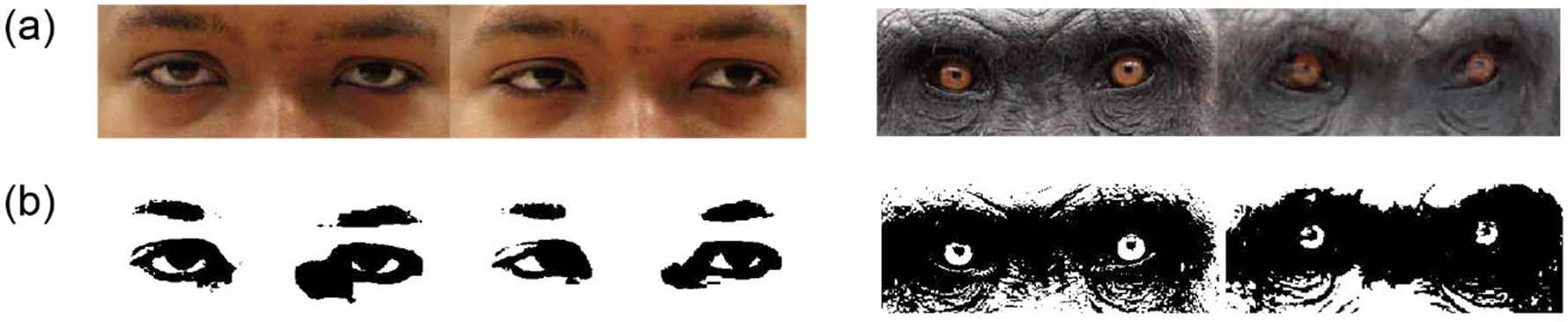
Examples of human (left) and chimpanzee eye images (right) in original forms (a) and in binarized forms (b) at an arbitrary brightness (LAB ΔE from black = 40).

In this study, we leveraged updated image analysis techniques to test the uniqueness of human external eye morphology among great apes to respond to these unanswered questions. Specifically, using the facial images of human and nonhuman great apes, we examined 1) the area-wise sclera exposedness by measuring the number of pixels in the iris and sclera, 2) the conspicuousness or saliency of eyes in the face using a visual saliency model, 3) the detectability of edges in eye and iris outlines using an edge-detection algorithm, and 4) the conspicuousness of eye outline and iris using a color-difference analysis. We examined both direct and averted gaze faces because many nonhuman individuals have pale colors in peripheral sclera areas, which are revealed only in averted gaze (Figure 1). Moreover, as the strength of a visual signal critically depends on natural noises such as darkening and distancing, we manipulated the brightness and blurriness (correlating with distance) of eye images and tested the detectability/conspicuousness of both eye outline and iris against these manipulations.

## 2. Material and methods

### Samples

We collected high-resolution images of seven great ape species, including bonobos (*Pan paniscus*), chimpanzees (*Pan troglodytes;* mostly *verus*, also including one *ellioti* and two hybrids), mountain gorillas (*Gorilla beringei beringei*), western lowland gorillas (referred to as ‘lowland gorillas’ in this paper; *Gorilla gorilla gorilla*), Bornean orangutans (*Pongo pygmaeus*), Sumatran orangutans (*Pongo abelii*), and humans (*Homo sapiens*). Nonhuman ape images were obtained through colleague researchers and keepers at zoos, research institutes, sanctuaries, and field sites (see Acknowledgements). Human images were obtained through The Columbia Gaze Data Set (Smith et al., 2013) which includes the faces of diverse ethnicities with various iris and skin colors.

From our collection of images, we selected 122 images of direct gaze faces (one image per individual) and 117 images of averted gaze faces (one image per individual). All images were of subadults or adults, and the number of females and males was balanced in each species (Table 1). We selected those images based on the following criteria: all parts of the faces were focused, zoomed, illuminated, and oriented toward the camera (we accepted deviations of ± 10 degrees for the direct gaze faces, and ± 45 degrees for the averted gaze faces) with relatively uniform lighting on the faces. Emotional expression was either absent or minimal in those faces. Each image was cropped to include central facial parts (eyes, nose, mouth and eyebrow ridges). The background was masked with a neutral color and not included in our analyses. The nonhuman ape images and human images were then converted into 345×460 and 400×400 pixels, respectively (this difference in width-height ratio was due to the difference in facial configurations). Finally, each image was auto-leveled in Photoshop to adjust tonal ranges within a face. See our online repository for the thumbnails of our images: https://osf.io/z6753/?view_only=5f6f393077194fb789c7fe5c44bbe06e

**Table 1.** The number of images/individuals (number of females in parentheses).

| Species | Direct gaze | Averted gaze | Total |
| --- | --- | --- | --- |
| *Pan paniscus* | 18 (13) | 17 (11) | 35 (24) |
| *Pan troglodytes* | 28 (14) | 29 (15) | 57 (29) |
| *Gorilla beringei* | 16 (9) | 14 (9) | 30 (18) |
| *Gorilla gorilla* | 11 (6) | 11 (8) | 22 (14) |
| *Pongo pygmaeus* | 10 (4) | 10 (4) | 20 (8) |
| *Pongo abelii* | 5 (3) | 2 (2) | 7 (5) |
| *Homo sapiens* | 34 (17) | 34 (17) | 68 (34) |
| Total | 122 (66) | 117 (66) | 239 (132) |

### Shape analysis

In each image, we traced outlines of eye-opening and iris with 2-pixel lines in Photoshop and then filled those traced lines to make a binary mask respectively for eye-opening and iris using a custom program in MATLAB v. 2019-2021 (MathWorks, Natick). We then measured 1) the maximal horizontal length (pixel) of the eye-opening mask, 2) the number of pixels in the eye-opening mask, and 3) the number of pixels in the sclera mask (by subtracting the iris mask and its outline from the eye-opening mask). Analogous to one-dimensional measures used in Kobayashi and Kohshima (2001), namely Sclera Size Index and Width/Height Ratio, we created the following two-dimensional measures: 1) Sclera Area Index, defined as the number of pixels in the sclera mask divided by the number of pixels in the eye-opening mask, and 2) Horizontal Elongation Index, defined as the squared maximal horizontal length of the eye-opening mask divided by the number of pixels in the eye-opening mask.

### Saliency analysis

To examine the conspicuousness of eyes in the images, we used a well-established saliency model (Itti & Koch, 2001) to calculate the pixel-level local saliency of the images (conspicuousness and saliency of color are exchangeable terms in this study). Although there are updated saliency models in the literature (see Sharma, 2015, for a review), we chose the Itti and Koch model implemented in MATLAB (Walther & Koch, 2006) due to its simplicity. This saliency model simulates primate visual attention and calculates local saliency within an image based on color (in the red-green and blue-yellow channels), intensity (lightness contrast) and orientation (the direction of edges/lines). The orientation information was discarded in this analysis because our purpose was to evaluate saliency of eye colors. We combined color and intensity information in the combined map with the weight of 1:1. To determine the saliency of the eye regions within each image, we defined the eye region as the rectangular area surrounding each eye and eyelid (see Figure 3a for an example) and calculated the sum of saliency values within the eye regions divided by the sum of saliency values within the whole image. We traced the background of each image and excluded them from the analysis. We also traced the pupils and any reflections of the light source in the eye of each image and excluded them from the analysis because the colors of these parts depend on the photographing environment but not on individuals/species.

**Figure 3.**
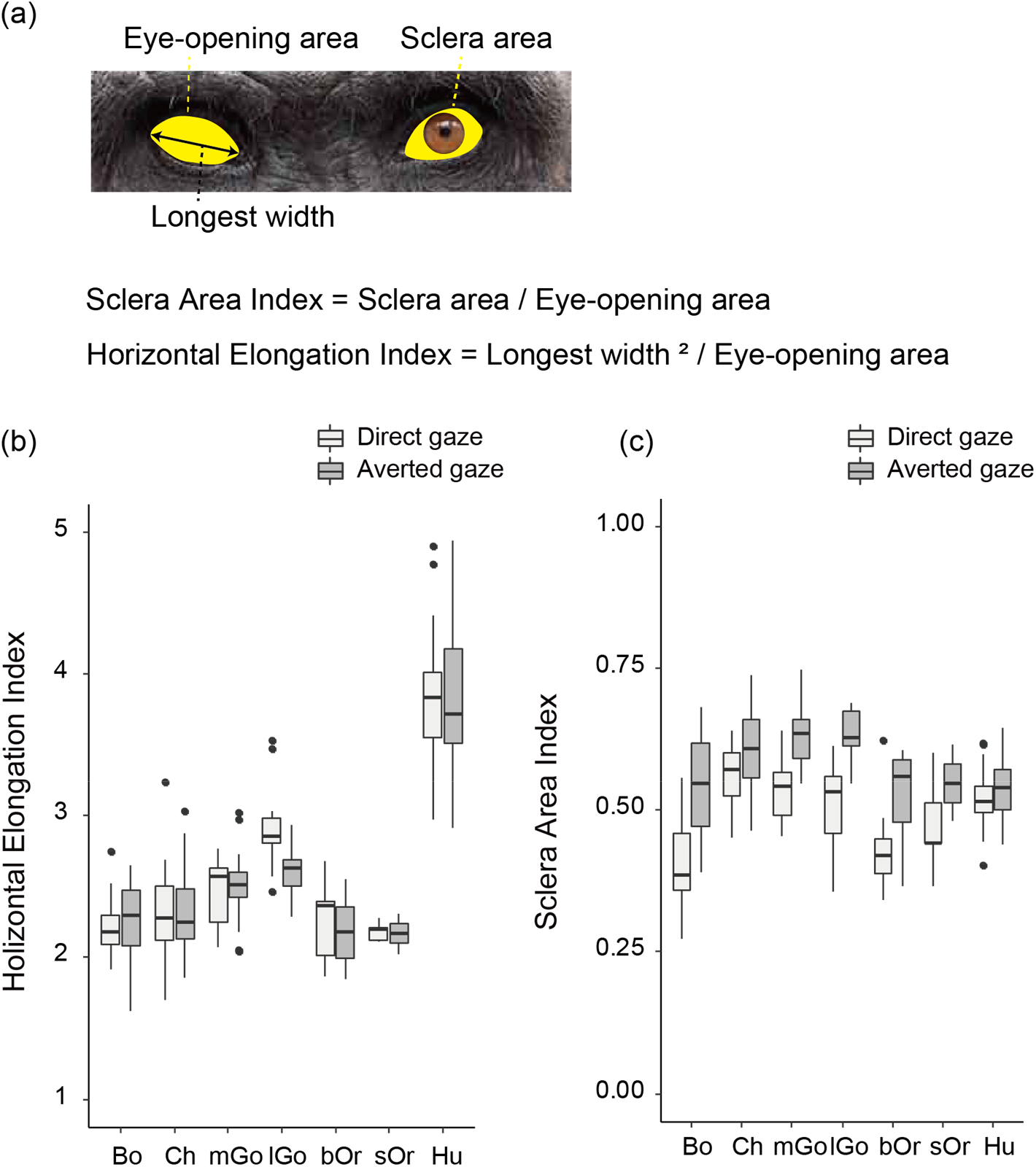
Eye shape of the great apes evaluated using our unique measures (a); Sclera Area Index (b) and Horizontal Elongation Index (c). Box plots show the median, interquartile range (IQR), and 1.5 × IQR, with outliers plotted individually. Bo = bonobos; Ch = chimpanzees; mGo = mountain gorillas; lGo = lowland gorillas, bOr = Bornean orangutans; sOr = Sumatran orangutans; Hu = humans.

### Edge detection analysis

To examine the visibility of iris and eye outlines in each image, we first converted RGB values of all pixels into brightness values (see below for the definition of brightness) in each image. We then ran an edge detection algorithm on those brightness-converted images using the ‘edge’ function with the ‘Sobel’ operator (default) in MATLAB. To simulate the visibility of detected edges at varying distances/blurriness, we blurred the original images (before the brightness conversion) using the ‘imgaussfilt’ function in MATLAB (the image processing toolbox) with a Gaussian of width, σ = 1, 2, 4, 8. This σ is the intensity of blur and is known to correspond linearly to the visual distances at which the image is presented (Coppens & van den Berg, 2004); i.e., the visual acuity measured with the image blurred with σ = 2 at 1 meter is identical to that measured with the image presented at 2 meters. To quantify the edges detected in the iris and eye outlines (and both combined) in each image (with σ = 1, 2, 4, 8), we created iris-outline and eye-outline masks by dilating those traced 2-pixel lines for each feature to 8 pixels in width and then calculated the proportion of pixels in which the edge was detected in these outline masks.

### Color analysis

To examine the color difference between eye features in each image, we used the CIE LAB color system. In this color system, L dimension represents lightness (0 to 100), A and B dimensions respectively represent red-green (−128 to 127) and blue-yellow (−128 to 127). The color difference was described as LAB ΔE, defined as Euclidean distance between the two LAB colors (i.e., the CIE 1976 style; although there are updated versions (e.g., CIE 2000), we used this older version for simplicity). Namely, LAB ΔE between color 1 and 2 was calculated as 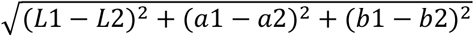. Throughout the paper, we refer to *lightness* as the lightness (or grayscale) component of a given color (i.e., L), *colorfulness* as chromatic components of a given color (i.e., 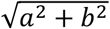, *brightness* as the color difference between a given color and black (L = 0, a = 0, b = 0; i.e., 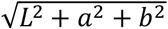, and *conspicuousness* as the color difference between a given color and its adjacent color(s).

To examine the color differences between each eye feature, we created outline and filled masks for iris, sclera, eye outline, and eye region using the rectangular eye region and 8-pixel outline masks described above. Those outline and filled masks did not overlap with one another. Namely, those filled masks did not include the 8-pixel outlines of iris and sclera outlines. The pupils and any reflections of the light source in the eye were also traced and excluded from the analysis. We then calculated the mean LAB of all pixels in each mask and the LAB ΔE between these means. We also measured the mean LAB of all pixels in the face region (the area excluding background) and covaried this value in the analysis to control for the variation in overall brightness of the whole face across individuals and species (this variation was mainly due to different skin colors across individuals and species and also partly due to the variations in lighting conditions across images; see our online repository). This control led to conservative (lower) estimation of color differences in our human samples (including diverse ethnicities) as their facial colors were generally brighter than those of other great apes.

To simulate the visibility of color differences between the masks at varying brightness (the color difference from black) levels of the images, we decreased the brightness of each image by dividing the RGB value of each pixel by a factor of 1, 2, 4, 8 (by definition, LAB ΔE decreases proportionally to this change). LAB ΔE around 1-3 is known as the ‘just noticeable difference’ in human perception (Stokes, Fairchild, & Berns, 1992).

Statistical analysis: We analyzed the images of direct and averted gaze faces separately because our aim was to test the uniqueness of human eyes in both gaze directions. To test species differences in eye properties, we used an analysis of variance (ANOVA) with species as between-subject factors in SPSS v. 23 (IBM Corp., 2015, Armonk). If ANOVA included a within-subject factor (e.g., different eye parts) and the assumption of sphericity was violated, a Greenhouse-Geisser’s correction was applied to the degrees of freedom. To directly test the uniqueness of eye properties in humans, we conducted follow-up tests comparing humans with each nonhuman species with the alpha level corrected for the number of pairwise comparisons in the Bonferroni correction. We excluded Sumatran orangutans from our analyses due to an insufficient number of samples (Table 1) but present their results in all graphs. We visually inspected the results from each Sumatran orangutan and confirmed that all individuals fit well within the individual/species variations of other nonhuman apes (see graphs). For the color analysis, we covaried the brightness of face mask in an analysis of covariance (ANCOVA).

## 3. Results

Shape analysis. We tested species difference in Sclera Area Index (the number of pixels in the sclera mask divided by the number of pixels in the eye-opening mask; Figure 3a), respectively for the direct and averted gaze faces (Figure 3b). ANOVA with species as a between-subject factor revealed significant main effects of species for both direct gaze faces (*F*[5, 111] = 17.65, *p* < 0.001, *η* _*p*_^2^ = 0.29) and averted gaze faces (*F*[5, 109] = 8.92, *p* < 0.001, *η*_*p*_^2^ = 0.44). Post-hoc pairwise comparisons between humans and the other species (*α* = 0.05/5) revealed that the sclera of humans is significantly more widely exposed than that of bonobos and Bornean orangutans (*p* < 0.001), less exposed than that of chimpanzees (*p* = 0.007), but not significantly different from that of mountain and lowland gorillas (*ps* > 0.3) in the direct gaze faces. In the averted gaze faces, the sclera of humans is significantly less exposed than that of chimpanzees, mountain gorillas, and lowland gorillas (*ps* < 0.001) but not significantly different from that of bonobos and Bornean orangutans (*ps* > 0.6).

We then tested species difference in Horizontal Elongation Index (the squared maximal horizontal length of the eye-opening divided by the number of pixels in the eye-opening mask; Figure 3a) respectively for the direct and averted gaze faces (Figure 3c). ANOVA with species as a between-subject factor revealed significant main effects of species for both direct gaze faces (*F*[5, 111] = 92.80, *p* < 0.001, *η*_*p*_^2^ = 0.81) and averted gaze faces (*F*[5, 109] = 73.32, *p* < 0.001, *η*_*p*_^2^ = 0.77). Post-hoc pairwise comparisons between humans and the other species (*α* = 0.05/5) revealed that the eye of humans is significantly more elongated than that of any other species in both direct (*ps* < 0.001) and averted gaze faces (*ps* < 0.001). These results indicate that the sclera of humans is not particularly more widely exposed than that of other apes, while the human eye is horizontally longer than other species’ eye.

### Saliency analysis

We tested species difference in eye saliency (the sum of saliency values within the eye region divided by the sum of all saliency values within the whole face) of the combined map respectively for the direct and averted gaze faces. An ANOVA with species as a between-subject factor revealed significant main effects of species for both direct gaze face (*F*[6, 109] = 3.99, *p* = 0.0021, *η*_*p*_^2^ = 0.16) and averted gaze faces (*F*[6, 111] = 9.93, *p* < 0.0001, *η* _*p*_ ^2^ = 0.31). Post-hoc pairwise comparisons between humans and the other species (*α* = 0.05/5) revealed that the eye of humans is significantly more salient than that of Bornean orangutans (*p* < 0.001), but not significantly different from that of other species (*ps* > 0.01) in the direct gaze faces, and significantly more salient than that of Bornean orangutans (*p* = 0.001) but not significantly different from that of other species (*ps* > 0.05) in the averted gaze faces. Figure 4b-c indicate that the saliency of chimpanzee and mountain gorilla eyes mainly depends on the colorfulness of their eyes (i.e., in the color map), while that of human eyes mainly depends on the lightness contrast of their eyes (i.e., in the intensity map). These results indicate that human eyes are not particularly more salient compared to other apes’ eyes.

**Figure 4.**
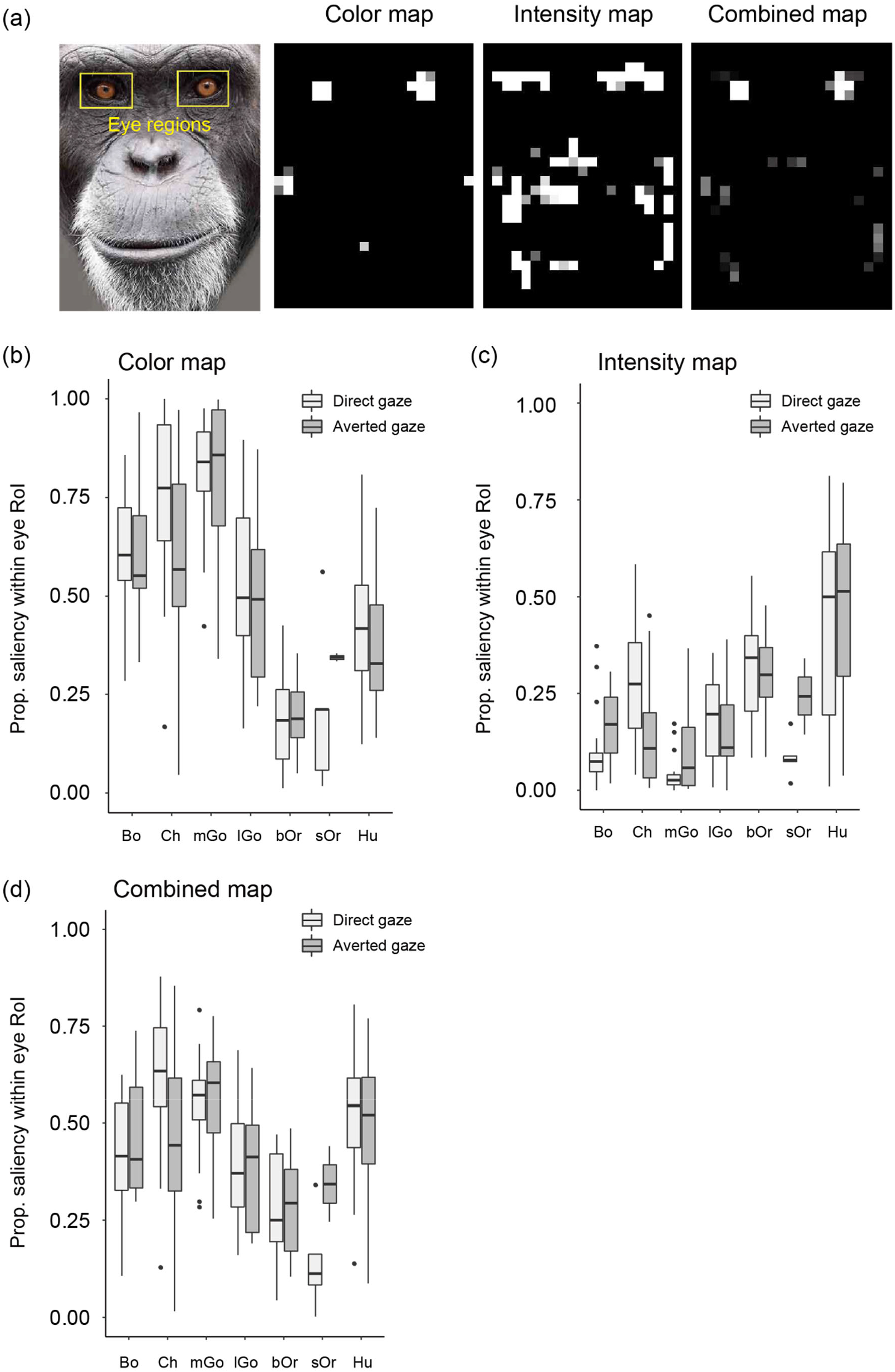
Saliency analysis. (a) Examples of eye region, the color, intensity, and combined map (in a chimpanzee face). (b) Proportion of pixels identified as salient in terms of colorfulness within the eye region, with respect to all pixels identified as salient within the whole image. (c) Proportion of pixels identified as salient in terms of intensity within the eye region, with respect to all pixels identified as salient within the whole image. (d) Proportion of pixels identified as salient in terms of both color and intensity within the eye region, with respect to all pixels identified as salient within the whole image. Box plots show the median, interquartile range (IQR), and 1.5 × IQR, with outliers plotted individually. Bo = bonobos; Ch = chimpanzees; mGo = mountain gorillas; lGo = lowland gorillas, bOr = Bornean orangutans; sOr = Sumatran orangutans; Hu = humans.

### Edge detection analysis

We tested species differences in detectability of edges on the combined mask for iris and eye outlines (the proportion of pixels in which edges were detected on the combined mask) on the brightness-converted images. To simulate visual distances, we manipulated the original images using Gaussian blurs with four levels of blur intensities (σ = 1, 2, 4, 8; Figure 5a). We then tested robustness of detected edges on the combined outline mask against such blurs across species (Figure 5d). A repeated-measures ANOVA with species as a between-subject factor and image blur as a within-subject factor revealed significant interaction effects between species and image blur for both direct gaze faces (*F*[8.8, 194.4] = 17.29, *p* < 0.001, *η*_*p*_^2^ = 0.44) and averted gaze faces (*F*[8.9, 193.6] = 21.16, *p* < 0.001, *η*_*p*_^2^ = 0.49). We then tested the species difference for the images blurred with σ = 8 using ANOVA with species as a between-subject factor. The main effects of species were significant for the images of both direct gaze faces (*F*[5, 111] = 18.96, *p* < 0.001, *η* _*p*_ ^2^ = 0.46) and averted gaze faces (*F*[5, 109] = 25.38, *p* < 0.001, *η* _*p*_ ^2^ = 0.54). Post-hoc pairwise comparisons between humans and the other ape species (α = 0.05/5) revealed that more edges were detected in the combined masks of humans than those of other species in both direct (*ps* < 0.001) and averted gaze faces (*ps* < 0.001). Similar patterns of results were observed respectively for iris-outline and eye-outline masks (Figure 5b-c). These results indicate that iris and eye outlines are more visible in human eyes than nonhuman ape eyes, particularly when the faces are blurred (or in a distant location).

**Figure 5.**
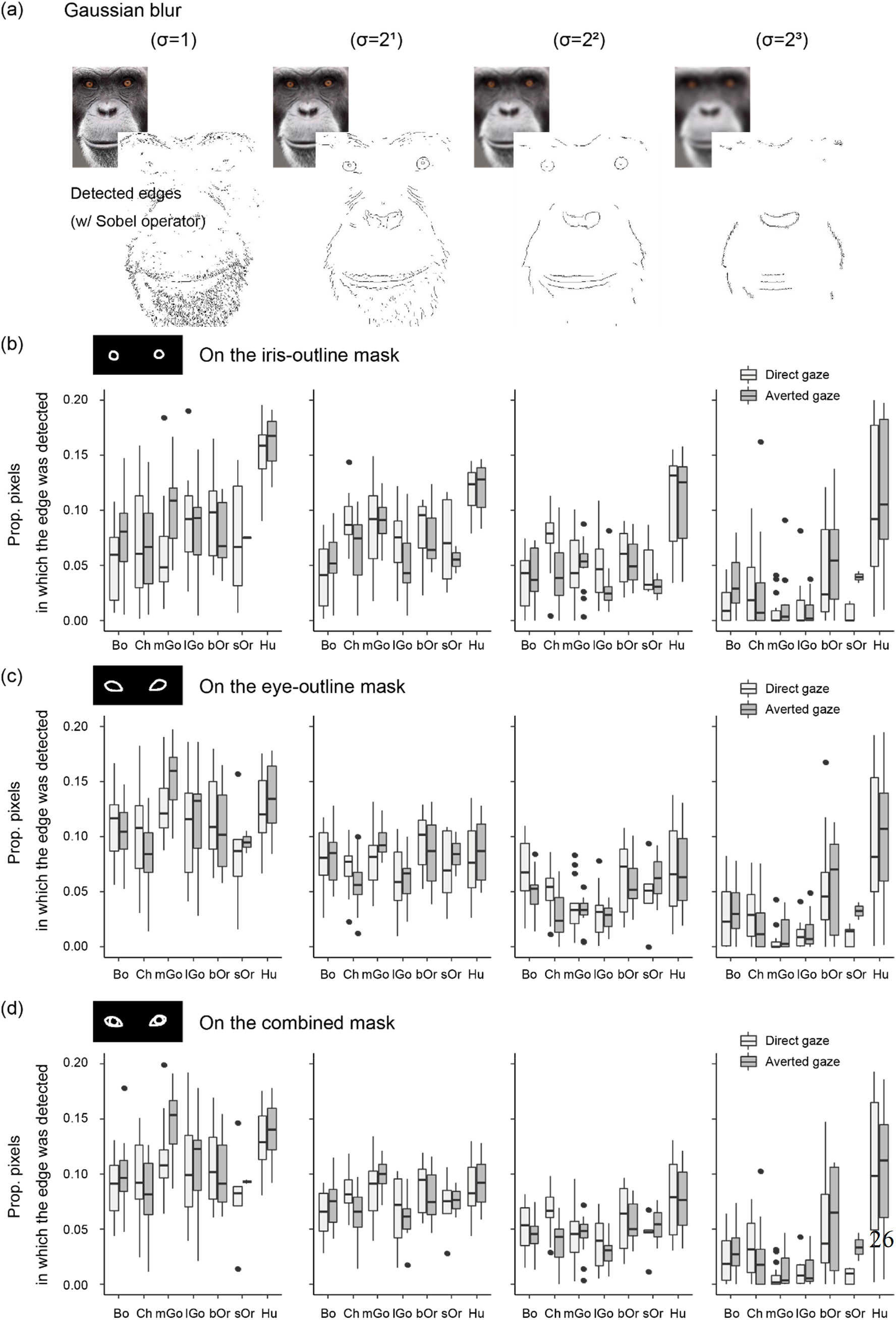
Edge-detection analysis and blurriness (or distance) manipulation test. Facial images were blurred with a Gaussian of width σ = 1, 2, 4, 8. (a) Examples of face images (top-left) and edges detected using an edge-detection algorism; b) Proportion of pixels in which the edge was detected on the iris-outline mask (with respect to all pixels on the iris-outline mask) as a function of image blur; c) Proportion of pixels in which the edge was detected on the eye-outline mask (with respect to all pixels on the eye-outline mask) as a function of image blur; d) Proportion of pixels in which the edge was detected on the combined mask (with respect to all pixels on the combined mask) as a function of image blur. Box plots show the median, interquartile range (IQR), and 1.5 × IQR, with outliers plotted individually. Bo = bonobos; Ch = chimpanzees; mGo = mountain gorillas; lGo = lowland gorillas, bOr = Bornean orangutans; sOr = Sumatran orangutans; Hu = humans.

### Color analysis

We tested species differences in conspicuousness of the combined mask for eye outline and iris (the color difference between the combined mask and elsewhere in the eye region; Figure 6g). An ANCOVA with species as a between-species factor and brightness in face region as a covariate revealed significant main effects in both direct (*F*[5, 110] = 22.87, *p* < 0.001, *η*_*p*_^2^ = 0.51) and averted gaze faces (*F*[5, 108] = 22.80, *p* < 0.001, *η*_*p*_^2^ = 0.51). Post-hoc pairwise comparisons between humans with the other species (α = 0.05/5) revealed that colors were more conspicuous in the combined mask of human eyes compared to that of any other species in both direct gaze faces (*ps* < 0.001) and averted gaze faces (*ps* < 0.001). Fig. 6a-f indicate that this result depends on the conspicuousness of both eye outline and iris in the human eye.

**Figure 6.**
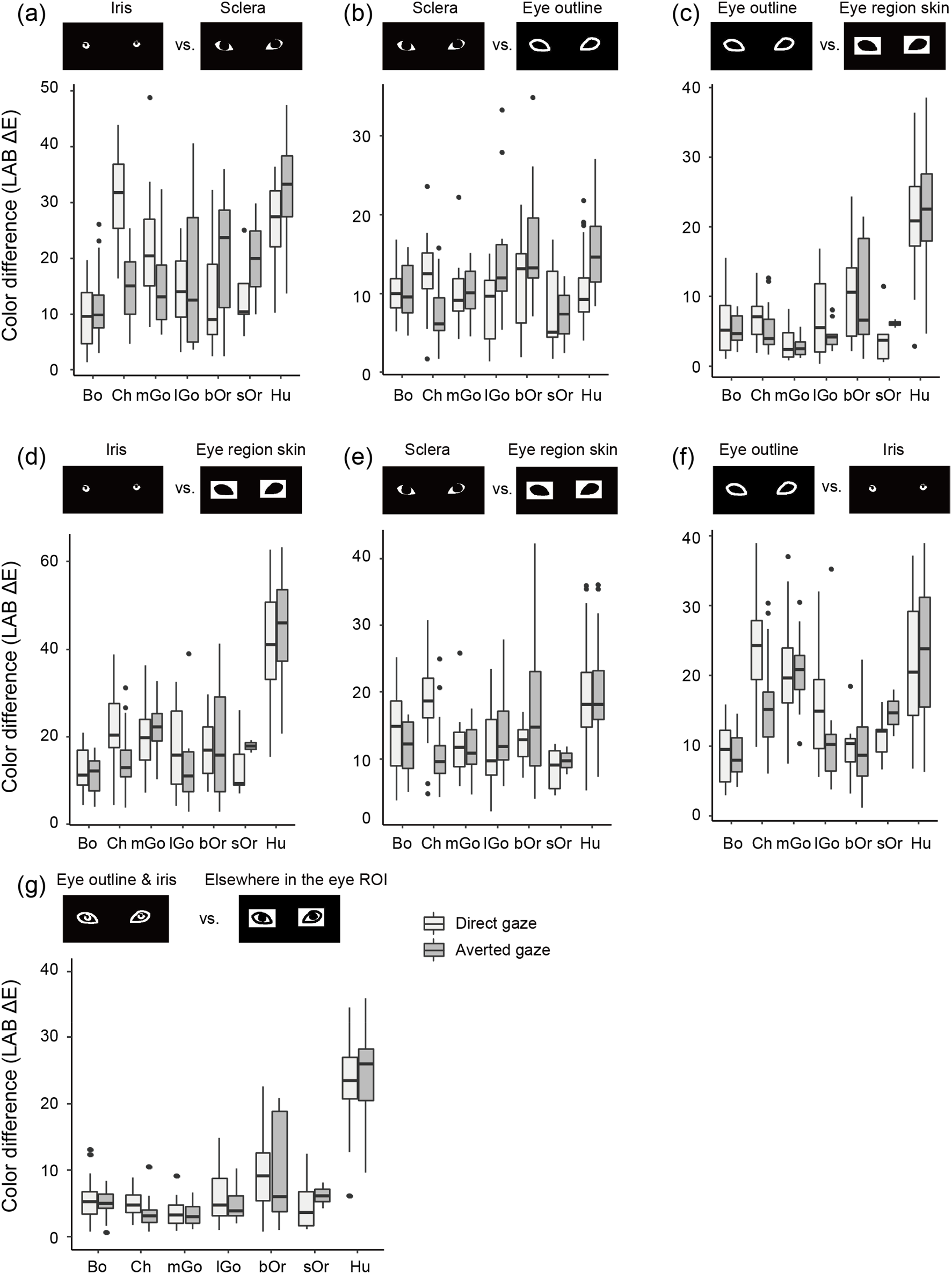
Color-difference analysis. (a-f) Pairwise color differences (LAB ΔE) between the iris, sclera, eye outline, and eye region skin masks. (g) Color differences (LAB ΔE) between the combined masks (the eye-outline and iris masks) and elsewhere in the eye region. Box plots show the median, interquartile range (IQR), and 1.5 × IQR, with outliers plotted individually. Bo = bonobos; Ch = chimpanzees; mGo = mountain gorillas; lGo = lowland gorillas, bOr = Bornean orangutans; sOr = Sumatran orangutans; Hu = humans.

Figure 7 presents how these color differences decrease as a function of image brightness (the color difference from black). We manipulated the image brightness by dividing the RGB value of each pixel by a factor of 1, 2, 4, 8. By definition, LAB ΔE decreased proportionally to this decrease in RGB. Given that LAB ΔE is barely noticeable around 1-3 in human perception (Stokes et al., 1992), the iris remains noticeable in the eye of most great ape species (including humans) in those dark images, while the eye outline remains noticeable only in the eye of humans in those dark images. Together, these results indicate that both iris and eye outlines are clearly visible in humans while eye outlines are particularly difficult to see in nonhuman apes, especially when the brightness of the face is low.

**Figure 7.**
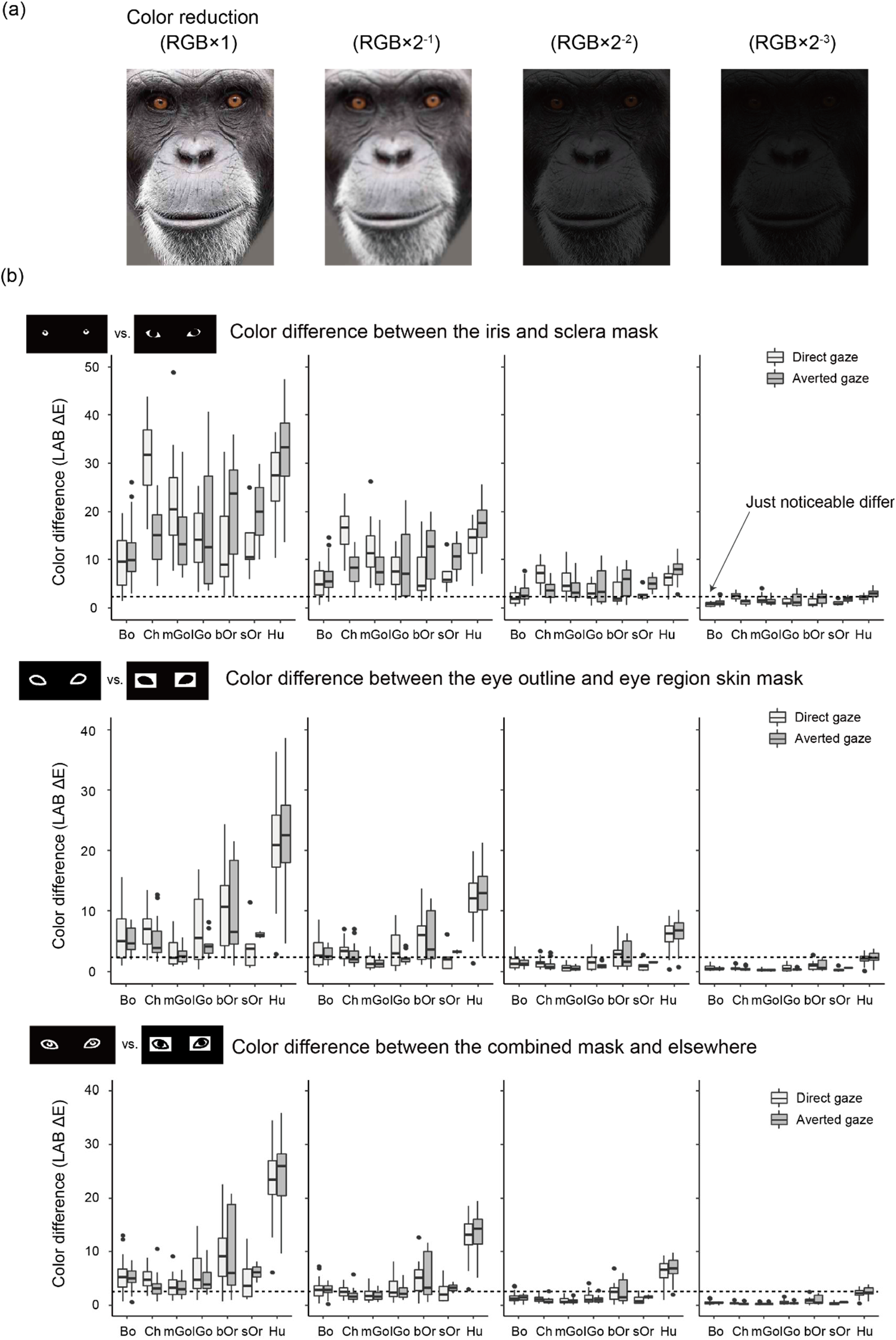
Color manipulation test. (a) Examples of a face image of which brightness was reduced (i.e., the RGB value of each pixel was divided) by a factor of 1, 2, 4, 8. (b) Color difference (LAB ΔE) between the iris and sclera masks, the eye outline and eye region skin masks, and the combined masks (the eye-outline and iris masks) and elsewhere in the eye region as a function of image brightness. The dotted line indicates the just noticeable difference (ΔE = ∼1-3) in human perception. Box plots show the median, interquartile range (IQR), and 1.5 × IQR, with outliers plotted individually. Bo = bonobos; Ch = chimpanzees; mGo = mountain gorillas; lGo = lowland gorillas, bOr = Bornean orangutans; sOr = Sumatran orangutans; Hu = humans.

## 3. Discussion

We found that the uniqueness of human eyes is characterized by clear visibility of both eye outline and iris, two essential features that critically contribute to the visibility of eye-gaze direction. These two unique features of human eyes are most likely attributable to the uniform whiteness in the exposed sclera, which makes both eye outline and iris particularly distinguishable, even in visually challenging conditions (e.g., at a distance or in poor lighting). Notably, eye outline is distinguishable in humans irrespective of skin color because eyelashes and shadows create thick dark lines surrounding each eye (see Figure 1 and 2). On the other hand, as the sclera color in great apes is either darker or more graded/patchy than that of human sclera, even though their iris is highly visible, their eye outline is less distinguishable from adjacent features, particularly when viewed in visually challenging conditions. Our results thus support a key premise of the gaze-signaling hypothesis, namely that human eyes are more distinguishable than those of nonhuman great apes. However, our results also challenge and critically update other key aspects of the gaze-signaling and game-camouflaging hypotheses. Specifically, 1) while the human eye is horizontally longer than the eye of other species, we have shown that the sclera of humans is not more widely exposed than that of other great ape species. Furthermore, we have demonstrated that 2) in both human and nonhuman apes, eyes are the most salient features of their faces and thus do not conceal gaze direction, and 3) the visibility of eye outlines critically depends on visual noise, such as a reduction in light or an increase in distance.

Regarding eye shape, our analyses identified that horizontal elongation is the only distinguished feature in the human eye. It could be argued that this feature may have evolved to signal eye-gaze direction particularly in the horizontal dimension. However, a key problem with this claim is that, as Kobayashi and Kohshima (2001) discovered, a horizontally elongated eye shape may have evolved to allow terrestrial primates to scan the environment more widely in the horizontal dimension. Thus, although it is possible that the more horizontally elongated eye of humans enhances the visibility of eye-gaze orientation at least in the horizontal dimension, it remains unclear whether this eye feature evolved for conspecific communication or other ecological reasons.

Regarding eye saliency, our analysis identified that eyes are salient features in the faces of all great ape species. Thus, consistent with Perea-García et al. (2019), this result questions the gaze-camouflaging hypothesis and instead suggests that great ape eyes are advertising rather than concealing their presence in the face. One might argue that the gaze-camouflaging hypothesis can be updated by proposing that the eye of nonhuman primates conceals eye-gaze direction rather than the presence of the eye *per se*. However, gaze direction can be inferred not only by eye orientations but also by head orientations (Emery, 2000). Moreover, nonhuman great apes tend to rely on head-directional cues rather than eye-directional cues to follow another’s gaze direction (Tomasello et al., 2007). Furthermore, our results showed that eye-gaze directions are reliably visible in both human and nonhuman great apes at least under good visual conditions. Thus, we believe that the gaze-camouflaging hypothesis should no longer be employed to describe the differences in eye color function between humans and the nonhuman great apes.

This study was the first to test the robustness of eye-gaze signal against visual noise by examinng the effect of image manipulations on the detectability/conspicuousness of eye outline and iris. One limitation of this approach is that we assumed similar visual perception across great ape species. We believe that this assumption is largely valid because previous electrophysiological, genetic, and behavioral studies identified similar color perception between humans and other apes, including habitual trichromatic vision (Deeb, Jorgensen, Battisti, Iwasaki, & Motulsky, 1994; Dulai, Bowmaker, Mollon, & Hunt, 1994; Jacobs & Deegan, 1999; Jacobs, Deegan, & Moran, 1996; Matsuno, Kawai, & Matsuzawa, 2004). However, slight differences might exist between humans and nonhuman apes in their spectrum sensitivity (Jacobs et al., 1996). Visual acuity and contrast sensitivity of nonhuman apes are similar or slightly inferior to those of humans (Adams, Wilkinson, & MacDonald, 2017; Bard, Street, McCrary, & Boothe, 1995; Matsuno & Tomonaga, 2006; Matsuzawa, 1990). Thus, despite reported similarities in visual perception between humans and nonhuman apes, experimental studies are essential to directly test the perceptual advantage of white sclera across species. Although several previous studies have shown that white sclera enhances gaze perception in humans (Ricciardelli, Baylis, & Driver, 2000; Yorzinski & Miller, 2020), it remains unclear whether this is due to special perceptual expertise of humans detecting white sclera (and a darker iris) as a result of experience, or instead to an intrinsic perceptual advantage provided by this morphological feature. Namely, it remains untested whether the perceptual advantage of white sclera can be observed even in a nonhuman species having a different iris-sclera color pattern from that of humans (i.e., dark sclera and a brighter iris in chimpanzees and mountain gorillas). Further studies are necessary in this respect.

In conclusion, our results support but also critically update the gaze-signaling hypotheses. Specifically, we found that a key unique characteristic of the human eye is clear visibility of both eye outline and iris, the two eye features that contribute to the visiblity of eye-gaze direction, rather than the extent to which sclera is exposed (area-wise) or the conspicuousness of eyes *per se* (in the face). Clear visibility of both eye outline and iris ensures the availability of eye-gaze signal in various visual conditions. Humans may employ such robust eye-gaze signal as a powerful communicative tool to leverage their everyday social interaction by constantly updating and exchanging information about their own and others’ attentional foci.

## Acknowledgements

We thank the following sanctuaries and zoos (in alphabetical order): Antwerp Zoo (especially, the Royal Zoological Society of Antwerp, Marjolein Osieck, Jeroen Stevens, Jonas Verhulst, and Sara Lafaut), Great Ape Research Institute (especially, Satoshi Hirata), Indianapolis Zoo, Kumamoto Sanctuary (especially, Toshifumi Udono, Etsuko Nogami, Melody So, and Lucy Baehren), Leipzig Zoo, Lincoln Park Zoo, Lola ya Bonobo, North Carolina Zoo (especially, Jennifer Ireland, Emily Lynch, Brooke Sides, and Chris Goldston), Olmense Zoo/Pakawi Park, Primate Research Institute (especially, Yoko Sakuraba and Yumi Yamanashi), Sepilok Orangutan Rehabilitation Centre (especially, Titol Malim, Sylvia Alsisto and Vijay S. Kumar), and Wolfgang Köhler Primate Research Center, and the following field sites: Wamba (especially Nahoko Tokuyama), Kalinzu, Bwindi, and Danum Valley (especially, Noko Kuze and Tomoko Kanamori) for kindly offering the images of great apes. We also thank the authors of Columbia Gaze Data Set for kindly offering the images of humans. We also thank Sabah Biodiversity Centre (License Ref. No. JKM/MBS.1000-2/2 JLD.11 (7) to T. Tajima) and Danum Valley Management Committee. Financial support came from Japan Society for the Promotion of Science KAKENHI Grants 19H01772 and 20H05000 to F.K, and the Lincoln Park Zoo Women’s Board supported J.G.L. and L.M.H.

## Notes

### Competing Interest Statement

The authors have declared no competing interest.

